# 42-parameter mass cytometry panel to assess cellular and functional phenotypes of leukocytes in bronchoalveolar lavage of Rhesus macaque

**DOI:** 10.1101/2024.09.19.613973

**Authors:** Mohau S. Makatsa, Anna Kus, Alice Wiedeman, S. Alice Long, Chetan Seshadri

**Affiliations:** Department of Medicine, University of Washington School of Medicine, Seattle, USA; Translational Immunology, Benaroya Research Institute, Seattle, WA, USA

**Keywords:** Mass cytometry, non-human primate, Immunophenotyping, Leukocytes, Bronchoalveolar lavage

## Abstract

This Optimized Multiparameter Immunofluorescence Panel (OMIP) reports on the development of a mass cytometry panel for broad immunophenotyping of leukocytes from bronchoalveolar lavage from rhesus macaques. Using this panel, we were able to identify myeloid populations such as macrophages, neutrophils, monocytes, myeloid and plasmacytoid DCs, basophils and lymphoid cell lineages including B cells, natural killer (NK) cells, mucosal associated invariant T (MAIT) cells, γδ T cells, CD4 T cells, CD8□β T cells, CD8 □□ T cells, and innate lymphoid cells (ILCs). We also included markers for defining memory, differentiation (CCR7, CD28, CD45RA), homing potential (CXCR3), cytotoxic potential (perforin, granzyme B, granzyme K), cell activation/differentiation (HLA-DR, CD69, IgD) and effector function (CD154, IFN-γ, TNF, IL-2, IL-17A, IL-6, IL-1β, CCL4 and CD107a). This panel was optimized on cryopreserved, bronchoalveolar lavage and splenocytes collected from rhesus macaques. The antibodies selected in this panel are human-specific antibodies that have been shown to cross-react with non-human primates except for CD45 clone D058-1283 which is specific for non-human primates.

## BRIEF NARATIVE

We developed this panel in our efforts to study correlates of vaccine-induced protection against *Mycobacterium tuberculosis* infection or TB disease in non-human primates (1). Since tuberculosis is primarily a disease of the lung, our goal was to broadly phenotype leukocytes in the bronchoalveolar lavage of rhesus macaques. Flow cytometry is widely used for immunophenotyping; however, due to the autofluorescence inherent to alveolar macrophages, it is difficult to characterize myeloid populations in the lung using fluorescence-based flow cytometry (2,3). Therefore, we employed mass cytometry, which uses metal conjugated rather than fluorescently-conjugated antibodies thus circumventing the autofluorescence from alveolar macrophages (4,5).

Myeloid cells can be distinguished using CD33 staining in humans (6); however, we did not find a CD33 antibody that cross-reacts with rhesus macaques. CD33 clone AC014.3E3 has been reported to react with rhesus macaque as listed on the NHP reagent resource reactivity database (NHP Reagent Resource), but it failed to stain any rhesus macaque cells from whole blood in our experiments (data not shown). Therefore, we did not include this marker to differentiate myeloid and lymphoid cells broadly. Instead, we used expression of surface CD68, CD206 and/or CD163 to distinguish macrophages and their subsets as shown in Figure 1a and Online Figure 1. We observed a low frequency of CD163 expressing macrophages, which is likely due to cryopreservation as studies have previously observed higher CD163 expressing macrophages in fresh bronchoalveolar lavage samples (1). Monocytes were identified by the expression of CD14 and CD11b expression with lack of macrophage markers (CD68, CD206, CD163). To identify granulocytes, we used anti-CD66abce antibody TET2 clone whose expression is limited to neutrophils and eosinophils. We used a combination of CD123, CD11c and HLA-DR to identify basophils (Lin-HLA-DR-CD123+), plasmacytoid dendritic cells (Lin-CD11c-HLA-DR+CD123+), and myeloid DCs (Lin-HLA-DR+CD11c+CD123-) while CD127 was used to identify Innate Lymphoid cells (Lin-CD127+). Lin-was defined as negative for T cells (CD3), B cells (CD20), macrophages (CD206, CD163, and/or CD68), monocytes (CD14, CD16), and neutrophils (CD66abce) and NK cells (NKG2A). We characterized myeloid function by expression of chemokine receptor (CCL4) and production of cytokines (TNF, IL-6 and IL-1β Fig. 1c).

**Figure 1.**
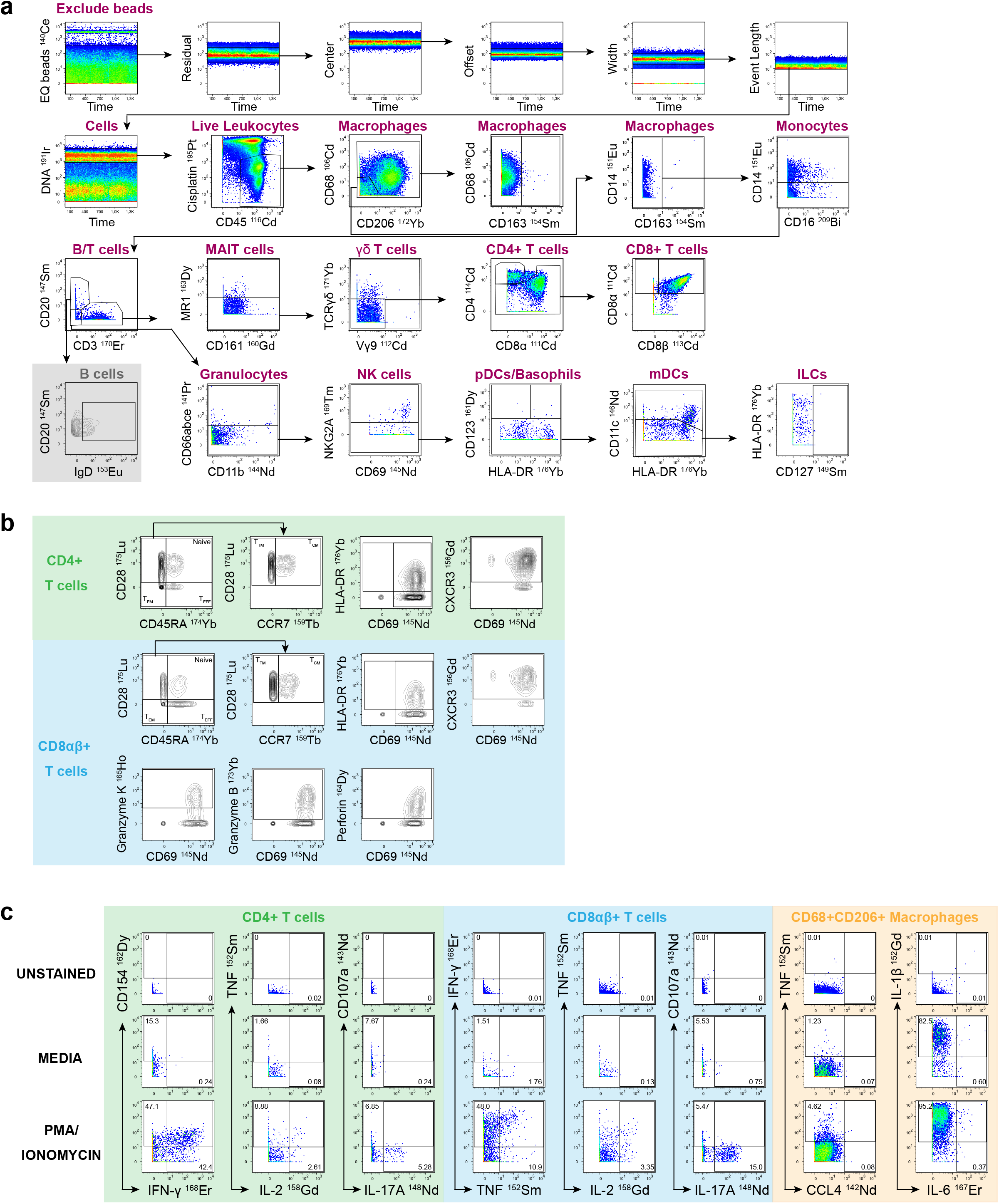
Immunophenotyping of non-human primate leukocytes subsets identified by mass cytometry in bronchoalveolar lavage using FlowJo manual gating. **(A)** EQ beads were first removed, and Gaussian discrimination parameters (Residual, Center, Offset, Width and Event Length) were then used to exclude debris, doublets, and aggregates. This was followed by gating on iridium DNA intercalator to identify nucleated cells, cisplatin to exclude dead cells, and CD45 to identify leukocytes. Macrophages were identified by CD68+, CD206+ and/or CD163+, monocytes by CD14+ and CD16 and lack of expression of macrophage markers. We used CD3 to identify T cells and CD20 to identify B cells. From the CD3-CD20-population, we gated Neutrophils by CD11b+CD66+, NK cells by NKG2A+, Basophils by HLA-DR-CD123+, plasmacytoid dendritic cells by CD11c-HLA-DR+CD123+, myeloid DCs by HLA-DR+CD11c+CD123-, and Innate Lymphoid cells by Lin-CD127+. B cells were further classified into two subsets based on IgD expression. Of the CD3+ cells, we further identified MAIT cells using MR1-5-O-PRU tetramer+, γδ T cells by Pan TCR γδ+, CD4+ T cells by CD4+, and CD8+ T cells were subdivided into CD8αβ (CD8α+CD8β+) and CD8αα (CD8α+CD8β-). **(B)** The memory phenotype of CD4+ and CD8+ T cells was determined based on the expression of CD45RA, CD28 and CCR7. Central memory (TCM) was classified as CD45RA-CD28+CCR7+, transitional memory (TTM) as CD45RA-CD28+CCR7-, effector memory (TEM) as CD45RA-CD28-and terminal effectors cells (TEF) as CD45RA+CD28-. CD8+ T cells were further characterized to determine activation profiles based on HLA-DR+ and/or CD69+, homing potential by CXCR3+ and cytotoxicity by granzyme B, granzyme K and perforin. This characterization can also be applied to other leukocytes subsets such as CD4, MAIT cells, γδ T cells or NK cells. **(C)** Chemokine and cytokine gating for CD4+ T cells, CD8+ T cells and macrophages. Gates were set from the unstained controls; unstimulated controls (media) were also included, and production of chemokines and cytokines were induced using Phorbol 12-myristate 13-acetate (PMA) and Ionomycin, which activate T cell intracellular signaling pathways. Functional makers that were used for lymphocytes are: CD154, IFN-γ, TNF, IL-2, IL-17A and CD107a while function makers that were used for macrophages are: TNF, IL-6, IL-1β, and CCL4. Although PMA/Ionomycin is not specific for myeloid stimulation, we were able to detect some myeloid chemokine (CCL4) and cytokine (IL-6, and TNF) signals above media control. IL-1β signal was detected without stimulation. Contour bivariate density plots were used for gating phenotypic characterization (memory profiles, activation, differentiation, homing and cytotoxic potential) because the topographical presentation of intensity in contour bivariate plots allowed easier discrimination of the positive from negative events. Pseudocolor bivariate dot plots were used for gating leukocytes cell subsets and functional properties where it was easy to distinguish the positive and negative events.

B cells can be identified using CD19 and/or CD20. In non-human primates, CD20 is predominantly used to identify B cells while CD19 is the classical marker for B cells in human studies (7,8). In this panel, we used CD20 for B cell identification and we included IgD, which is typically used together with CD27 to distinguish between naïve and memory B cells (9,10; Fig. 1a).

For identification of NK cells, we used NKG2A on CD3-cells. NKG2A is widely used as the marker for NK cells in non-human primates (11). We did not classify the NK “like” T cells (NKG2A+CD3+) because NKG2A can also be expressed by a subset of CD8+ T cells with effector phenotype (16). However, if annotation of NK “like” T cells is required, these cells can be identified as CD3+CD8α-NKG2A+. Using unsupervised clustering methods may enable distinct identification of NK “like” T cells in non-human primates. Notably, these cells are not annotated as NKT cells, which are defined by the expression of a semi-invariant T cell receptor (12).

This panel allowed for identification of donor-unrestricted T cells by using MR1 5-OP-RU tetramer to identify Mucosal-associated invariant T cells (MAIT) and Pan TCR γδ+ and TCR Vγ9 antibodies to identify γδ T cells from the CD3+ leukocytes (13; Fig. 1a). We also identified conventional T cells by their expression of CD4 and CD8 (Fig. 1a). CD8 molecule has been shown to express two CD8 isoforms that pair on the cell surface as either a CD8αα-homodimer or as a CD8αβ-heterodimer (14,15). CD8β is more conserved and mostly restricted to CD8 T cells while CD8α has been shown to be expressed by other cell subsets, including NK cells, MAIT cells, and γδ T cells (16). In this panel, we included both CD8α and CD8β and we were able to identify both CD8αβ+ and CD8αα+ T cells. We also included markers to determine the cytotoxic potential (granzyme B, granzyme K and perforin; Fig. 1b) of NK cells, γδ, and CD8+ T cells (Fig. 1b).

CD4 and CD8 T cells were further characterized based on memory profiles using CD45RA, CD28 and CCR7 to classify central memory (CD45RA-CD28+CCR7+), transitional memory (CD45RA-CD28+CCR7-), effector memory (CD45RA-CD28) and terminal effector T cells (CD45RA+CD28-). Additionally, we examined the activation profiles based on CD69 and HLA-DR expression and homing potential predicted by CXCR3 expression (Fig. 1b). Lastly, we determined functional characteristics of leukocyte subsets, using CD154, IFN-γ, TNF, IL-2, IL-17A and CD107a (Fig. 1c).

To summarize, we developed a 42-parameter mass cytometry panel (including viability by cisplatin and DNA contentment by iridium) to enable the identification and characterization of leukocyte populations in bronchoalveolar lavage of rhesus macaques (Table 1). Data analysis can be accomplished using manual gating in FlowJo or by using unsupervised clustering methods such as that developed by Norwicka et al, (17) which uses FlowSOM clustering to define cell populations that can be manually annotated using canonical markers by an expert (Online Fig. 1).

**TABLE 1.** Summary table for application of OMIP-.

|  |  |
| --- | --- |
| Purpose | Immunophenotyping of leukocytes in bronchoalveolar lavage of rhesus macaques including myeloid (macrophages, monocytes, neutrophils, mDCs, pDCs, and basophils) and lymphoid cell lineages (ILCs, B cells, NK cells, MAIT cells, $\gamma\delta$ T cells, CD4+ T cells and CD8+ T cells); characterization of memory differentiation, activation profiles and homing potential as well as cytotoxicity and functional properties. |
| Species | rhesus macaques |
| Cell Type | Cryopreserved bronchoalveolar lavage |
| Cross-References | Mass cytometry panels OMIP-088, -087, -054, -048, -045, and -034; panels that include human myeloid phenotyping OMIP-101, -062, -051, and -042 and rhesus macaque panels OMIP-052, -035, -029, -026, -016, and -005. |

**TABLE 2.**
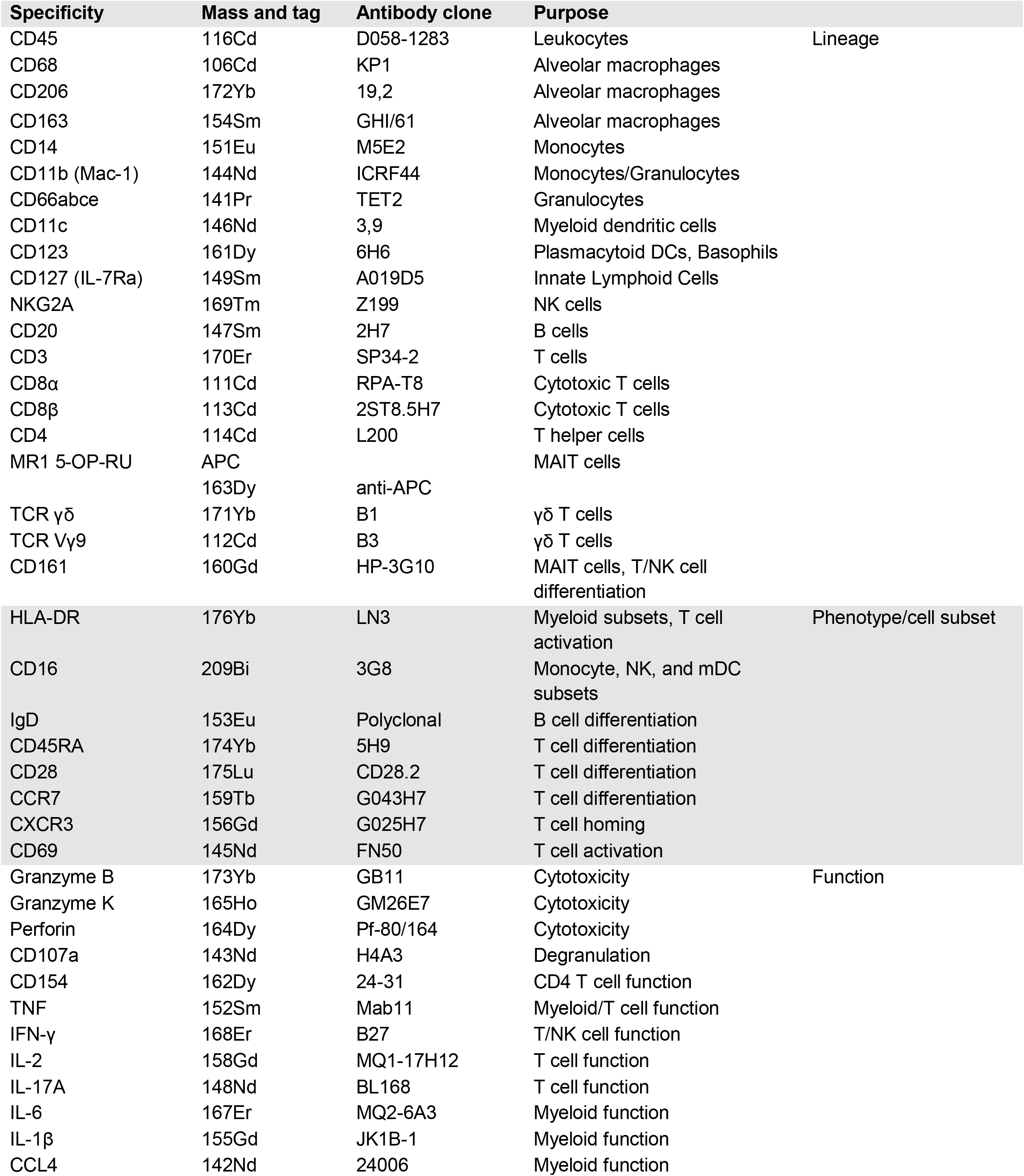
Reagents used in OMIP.

## SIMILARITY TO RELATED OMIPS

To our knowledge, this mass cytometry panel is the first to identify and characterize broad leukocytes subsets in the bronchoalveolar lavage of rhesus macaques. Although OMIP-088, -087, -054, -048, - 045, and -034 are mass cytometry panels, these focus on human peripheral leukocytes (OMIP-034), human head and neck tumors and cancer cell lines (OMIP-045), murine lymphocytes from spleen and lymph nodes (OMIP-048), mouse brain cells (OMIP-054), human peripheral blood mononuclear cells (OMIP-087), and mouse liver tissue (OMIP-088); and none of them focus on myeloid phenotyping of rhesus macaque. OMIP-101, -062, -051, and -042 include some of the myeloid markers used in this panel but these were focused on human cells and only OMIP-62 analyzed cells from the airway. Furthermore, OMIP-101, -062, -051, and -042 are flow cytometry panels and use fluorophore tagged rather than heavy metal tagged to identify leukocyte subsets.

## ACKNOWLEDGEMENTS

The authors would like to thank the Washington National Primate Research Center for providing bronchoalveolar lavage, whole blood and splenocytes samples used for initial panel optimization. Funding Statement: NIH Contract 75N93019C00071, P30-AI168034, and R01-AI146072 to C.S.

## AUTHOR CONTRIBUTIONS

Mohau S. Makatsa: Conceptualization; data curation; formal analysis; methodology; visualization; validation; writing – original draft; writing – review and editing.

Anna Kus: Conceptualization; methodology; writing – review and editing. Alice Wiedeman: Supervision; writing – review and editing.

S. Alice Long: Conceptualization; resources; supervision; writing – review and editing.

Chetan Seshadri: Conceptualization; funding acquisition; resources; supervision; writing – review and editing.

