## Supplemental material for "42-parameter mass cytometry panel to assess cellular and functional phenotypes of leukocytes in bronchoalveolar lavage of Rhesus macaque"

– ONLINE MATERIAL –

### PANEL DEVELOPMENT STRATEGY

The objective for developing this OMIP was to comprehensively profile leukocytes in the bronchoalveolar lavage of rhesus macaques. While flow cytometry is commonly used for immunophenotyping, the inherent autofluorescence of alveolar macrophages poses challenges in characterizing myeloid populations accurately (1). To overcome this issue, we employed a high-throughput mass cytometry assay, utilizing metal-conjugated antibodies to analyze cellular composition and characteristics of cells, thereby overcoming autofluorescence interference (2). Unlike Flow cytometry, mass cytometry exhibits minimal and predictable spill-over, which may arise from metal isotope impurity leading to ion spreading into adjacent channels, abundance sensitivity spreading, or oxidation events (3).

To optimize this panel, we first identified and screened non-human primate reactive antibodies by assigning each marker to a metal isotope. Highly expressed markers were assigned to lower signal isotopes and lowly expressed markers assigned to higher signal isotopes. We also used Maxpar Panel Designer to estimate the signal and tolerance values for assigned metal isotopes. Commercially available antibodies were obtained from Standard BioTools (formerly Fluidigm). All other antibodies were conjugated in house by labeling purified antibodies with the appropriate Maxpar Antibody Labeling Kit (Standard BioTools) according to the manufacturer's protocol. The Maxpar X8 Antibody Labeling Kit was used for conjugating antibodies to lanthanide metal isotopes (CD66abce 141Pr, CCL4 142Nd, CD107a 143Nd, CD69 145Nd, IgD 153Eu, IL-1β 155Gd, CD161 160Gd, CD123 161Dy, CD154 162Dy, Perforin 164Dy, Granzyme K 165Ho, IL-6 167Er, TCR γδ 171Yb, CD206 172Yb, CD45RA 174Yb, CD28 175Lu and HLA-DR 176Yb), and the Maxpar MCP9 Antibody Labeling Kit was used for conjugating antibodies to cadmium metal isotopes (CD45 116Cd, CD68 106Cd, CD4 114Cd, CD8α 111Cd, TCR Vγ9 112Cd, and CD8β 113Cd). We then performed serial two-fold dilutions and chose the optimal concentration for each antibody conjugate based on resolution of the positive population from the negative population and the frequency of the positive events (Online Fig. 2, Online Table 2).

Previous reports using flow cytometry suggest that CD56 is not a suitable marker for identification of NK cells in non-human primates, and NKG2A/C, CD16 and CD8α expression on CD3 negative cells have been suggested as more accurate markers for NK cell identification (4,5). Consistent with these reports, we show that NKG2A and CD56 co-staining of splenocytes from rhesus macaques mostly identify distinct cells with only 0.13% of CD3- leukocytes expressing both NKG2A and CD56 (Online Fig. 3). Almost all the NKG2A positive cells expressed CD8α (99%), while only 58% of CD56 positive cells were also positive for CD8α. Moreover, we observed higher expression of CD14 on CD56+CD3- leukocytes compared to NKG2A+CD3- leukocytes (66%

and 21%; respectively). In contrast to CD56+CD3- cells, NKG2A+ CD3+ cells were associated with higher expression of CD16 (43%) and higher cytotoxicity based on perforin expression (87%). These CyTOF data support previous flow cytometry reports highlighting NKG2A as a more suitable marker for identification of NK cells in non-human primates.

After assigning metal isotopes to each antibody, we performed a metal minus multiple (MMM) experiment to assess the potential for spillover in our panel. This entailed strategically dividing antibodies into three panels such that channels in which spillover resulting from abundance sensitivity, isotopic impurity, or oxidation would manifest were empty/open (Online Table 3). This allowed us to investigate if the stained antibody (mass) is spilling over into adjacent channels (mass-1 and mass+1) or in the oxidation channel (mass+16). During the panel development, we had initially assigned HLA-DR to 153Eu. However, our MMM experiment revealed a spillover of HLA-DR 153Eu into CD14 151Eu which prevented accurate identification of CD14 expressing cells (Online Fig. 4). Online Figure 4a shows a spillover signal in the mass-2 channel (CD14 151Eu) after staining with HLA-DR 153Eu. True CD14 151Eu signal was not distinguishable above the contaminating signal from HLA-DR 153Eu (Online Fig. 4b). Therefore, HLA-DR was re-assigned to 176Yb which limits the spillover because there is no mass+1 (channel 177) or mass+16 (channel 192). We performed another MMM for the final panel (Online Table 3 and Online Fig. 5). We utilized tSNE high-dimensionality reduction to determine the presence of signal due to spillover by assessing the scaled marker expression of each stained antibody in its own detection channel (mass) as well as in mass-1, mass+1, and mass+16 channels. As previously mentioned, the MMM experiment is designed such that the mass-1, mass+1, and mass+16 channels are left open, so any signal detected in channels outside of a stained antibody's detection channel indicates spillover. Using this strategy, we identified signal from CD206 172Yb spilling into the adjacent mass-1 channel, which captures signal from granzyme B 173Yb. However, since granzyme B expression is characterized on CD206 negative cells, we concluded that this spillover would not affect data analysis. Similarly, we observed HLA-DR 176Yb signal spilling into CD28 175Lu (mass-1) and CD45RA 174Yb (mass-2) as a result of abundance sensitivity and metal impurity. Based on the location of the HLA-DR 176Yb signal on the tSNE plot, we noted that the spillover signal was limited to non-T cells (CD3-; Online Fig. 5), therefore this spillover signal does not affect CD45RA and CD28 analysis which are profiled on T cells.

**DATA ACQUISITION, NORMALIZATION AND ANALYSIS**

Immediately before acquisition, cells were washed with ultrapure water and resuspended in a solution of EQ Four Element Calibration Beads (Standard BioTools) diluted 5-fold into cold ultrapure Water and filtered through cell strainer cap tubes (Fisher Scientific). Samples were then acquired on a CyTOF Helios instrument (Standard Biotools. Online Table 1) at an event rate no greater than 500 events per second. FCS files were normalized for EQ Bead intensity using the CyTOF Software v 7.0.8493, as previously described (6). Normalized fcs files were used for analysis in FlowJo 10.8.1. We manually gated to exclude EQ Beads, debris, doublets and ion cloud fusions using Gaussian discrimination parameters (Residual, Center, Offset, Width and Event Length; Figure 1; (7)). We then used DNA 191Ir to identify nucleated cells, Cisplatin to identify live cells, and CD45 116Cd to identify leukocytes. Further analysis on CD45+ leukocytes was conducted using bimodal gating on FlowJo v10.8.1 (Fig. 1), and unsupervised gating using the tSNE FlowJo Plugin (Online Fig. 4a and Online Fig. 5) and FlowSOM (Online Fig. 1) using a previously reported R workflow (8,9).

**CROSS REFERENCE:** There currently no panels that use mass cytometry to analyze broncho alveolar lavage samples from rhesus macaque, but there are panels that include at least one aspect described in this OMIP such as panels that employ mass cytometry: OMIP-088, -087, -054, -048, -045, and -034; panels that include human myeloid phenotyping OMIP-101, -062, -051, and -042 and rhesus macaque panels OMIP-052, -035, -029, -026, -016, and -005.

### RHESUS MACAQUE SAMPLES

Indian-origin rhesus macaques (Macaca mulatta) were used in this study. Macaques were housed and cared for in accordance with local, state, federal, and institute policies in facilities accredited by the American Association for Accreditation of Laboratory Animal Care (AAALAC), under standards established in the Animal Welfare Act and the Guide for the Care and Use of Laboratory Animals. Macaques were monitored for physical health, food consumption, body weight, temperature, complete blood counts, and serum chemistries.

### BRONCHOALVEOLAR LAVAGE PROCESSING

Tubes containing BAL were centrifuged at 300x*g* for 10 minutes to pellet cells and supernatant aspirated. The pellet was resuspended in 10 mL of cRPMI (10% Fetal bovine serum (FBS), 10 U/mL Penicillin/streptomycin in RPMI 1640 medium) with Benzonase (20 µL/10mL) and passed through a 70 µm filter. BAL cells were centrifuged again at 300x*g* for 10 minutes and supernatant aspirated, then frozen down in 10% Dimethyl sulfoxide (DMSO) in FBS at a concentration of approximately 1-3 million cells/ml.

**STAINING PROTOCOL:**

*Materials and reagents*

Falcon™ Round-Bottom Polystyrene Test Tubes with Cell Strainer Snap Cap (Thermo Fisher Scientific)

Nunc Nunclon D MicroWell Plates; 96-well U-bottom (Thermo Fisher Scientific)

Nunc™ 50mL Conical Sterile Polypropylene Centrifuge Tubes (Thermo Fisher Scientific)

Dulbecco's Phosphate-Buffered Saline (DPBS; Fisher Scientific)

Cytiva Hyclone™ Defined Fetal Bovine Serum (FBS; Fisher Scientific)

Penicillin-Streptomycin (10,000 U/mL; Thermo Fisher Scientific)

RPMI 1640 Medium (Fisher Scientific)

Bovine serum albumin (BSA; Sigma Aldrich)

Maxpar Cell Staining Buffer (CSB; Standard Biotools)

Maxpar Fix I buffer (Standard Biotools)

Maxpar Perm-S buffer (Standard Biotools)

Maxpar Fix and Perm Buffer (Standard Biotools)

EQ Four Element Calibration Beads (Standard Biotools)

Cell-ID Intercalator-Ir (Standard Biotools)

Paraformaldehyde (Sigma Aldrich)

Phorbol-12-myristate-13-acetate (PMA; Sigma Aldrich)

Ionomycin (Sigma Aldrich)

Brefeldin A (Sigma Aldrich)

BD GolgiStop Protein Transport Inhibitor (Fisher Scientific)

Tryphan blue (Sigma Aldrich)

*Thawing*

1. Add 10 mL warm complete RPMI (cRPMI, RPMI 1640 Medium with 10% Fetal Bovine Serum and 10 units/mL Penicillin-Streptomycin) and 20 uL Benzonase to a 50 mL conical tube for each sample.
2. Pull samples from the liquid nitrogen tank and thaw in a 37 °C water bath.
3. When a small (pea-sized) bit of ice remains in the cryovials, transfer the cryovials to the biosafety cabinet. *Dry off the outside of the cryovials, and wipe with alcohol solution before opening.*
4. Using a p1000 pipet, remove 1 mL of thaw media from the stock tube and add to each cryovial by dripping slowly down the inner side of the cryovial.
5. Transfer the cells from the cryovial to the 50 mL conical tube containing Benzonase and 10 mL warm cRPMI for each respective sample.
6. After transfer, rinse the cryovial once using 1 mL of thaw media and transfer to the 50 mL conical.
7. Centrifuge the conical tubes at room temperature at 500x*g* for 5 minutes.
8. Add 5 mL of pre-warmed cRPMI to each tube.
9. Count cells manually using Tryphan blue.
10. Resuspend cells in 50 mL conical at 2-5 million cells/mL for 2 hours in a 37 °C incubator with 5% $CO_2$.

*Stimulation*

11. Transfer up to 3 million cells/per well into a U-bottom tissue culture plate in 100 µL.
12. Add 2 µL PMA and 2 µL Ionomycin (final concentrations 50 ng/mL and 500 ng/mL; respectively)
13. Add 2 µL CD107a 143Nd and fill with cRPMI to a total of 200 µL per well.
14. Incubate plate in a 37 °C incubator with 5% $CO_2$ for a total of 18 hours.
15. At 12 hours of the stimulation, add 20 µL Brefeldin A (final concentration: 10 µg/mL) and 1x GolgiStop Protein Transport Inhibitor (1.6 µL).
16. After 18 hours of stimulation, proceed with staining immediately.

*Viability Staining*

17. Centrifuge cells at 500x*g* for 5 minutes.
18. Wash cells twice with 200 µL PBS and centrifuge at 500x*g* for 5 minutes.
19. Resuspend cells in 50 uL Cisplatin 195Pt to a final concentration 1 µM.
20. Incubate for 5 minutes at room temperature.
21. Quench the Cisplatin reaction with addition of 150 µL FACS buffer (0.2% BSA in PBS) to each well.
22. Centrifuge at 500x*g* for 5 minutes.
23. Wash twice with 200 µL FACS buffer and centrifuge at 500x*g* for 5 minutes.

*MR1 tetramer staining*

24. Add 50 uL MR1 5-OP-RU tetramer mix in FACS buffer.
25. Incubate for 60 minutes at room temperature.
26. Add 150 µL Maxpar Cell Staining Buffer (CSB) and centrifuge at 500x*g* for 5 minutes.
27. Wash twice with 200 µL CSB and centrifuge at 500x*g* for 5 minutes.

*Surface staining*

28. Add 50 µL surface marker antibody mix to cells and incubate for 30 minutes at 4°C.
29. Add 150 µL CSB and centrifuge at 500x*g* for 5 minutes.
30. Wash twice with 200 µL CSB and centrifuge at 500x*g* for 5 minutes.

*Intracellular staining*

31. Add 100 µL of 1x Maxpar Fix I buffer (1 in 5 dilutions in water) and mix.
32. Incubate for 20 minutes at room temperature.
33. Wash by adding 100 µL Maxpar Perm-S buffer and centrifuge at 700x*g* for 3 minutes.
34. Wash again with 200 µL Maxpar Perm-S buffer and centrifuge at 700x*g* for 3 minutes.
35. Add 50 µL intracellular marker antibody mix to cells and incubate for 30 minutes at 4°C.
36. Add 150 µL Maxpar Perm-S buffer and centrifuge at 700x*g* for 3 minutes.
37. Wash twice with 200 µL Maxpar Perm-S buffer and centrifuge at 700x*g* for 3 minutes.
38. Resuspend cells in 100 µL of 1% paraformaldehyde for 15 minutes at 4°C. *1% paraformaldehyde prepared fresh every time.*
39. Add 100 µL Maxpar Perm-S buffer and centrifuge at 700x*g* for 3 minutes.

*DNA intercalator staining*

40. Resuspend samples in 100 µL of 0.5 µM Cell-ID Intercalator-Ir solution prepared in Maxpar Fix and Perm Buffer.
41. Store samples at 4°C in the cell intercalation solution until time of acquisition. Samples stored in cell intercalation solution for up to 1 week at 4°C prior to acquisition (10). Longer storage of stained samples at -80 °C is also possible (10).

*Acquisition*

42. On day of acquisition, remove samples from 4°C and centrifuge at 700x*g* for 3 minutes.
43. Wash cells by adding 200 µL of CSB and centrifuge 700x*g* for 5 minutes.
44. Wash with 200 µL of cold ultrapure water and centrifuge 700x*g* for 5 minutes.
45. Resuspend cells in 200 µL solution of cold EQ Four Element Calibration Beads diluted 5-fold in ultrapure water.
46. Filter cells through Falcon™ Round-Bottom Polystyrene Test Tubes with Cell Strainer Snap Cap 35µM.
47. Adjust cell concentration to 0.5 x $10^6$ cells/mL in cold 1/5th EQ Four Element Calibration Bead solution. Filtered cells kept on ice until acquisition on instrument.
48. Acquire cells on CyTOF Helios Mass Cytometer at an acquisition rate no greater than 500 events per second.

### Online Table 1. Instrument Configuration

| | |
|---|---|
| Instrument | Helios Mass Cytometer (Standard Biotools) |
| CyTOF Software | Version 7.0.8493 |
| Flow rate | 30µl/min |
| Cell concentration | $0.5 \times 10^6$ cell/ml |
| Sample carrier | 1/5 EQ beads in ultrapure water |
| Event rate | 250-500 events/s |
| Lower convolution threshold | 400 |
| Acquisition settings | Noise reduction used |
| Pre-processing of data | Data normalization using internally spiked EQ beads (1/10 v/v) using Standard Biotools software |
| Tuning frequency | Daily |
| Tuning results range (Tb) | 763,878-1,607,567 Dual Counts |
| EQ Bead Passport | EQ-P13H2302_ver2 |
| Min and Max Event Duration | 10-150 |

**Online Table 2. Summary table for antibodies used in the OMIP.**

| Specificity | Mass/tag | Clone | Company | Catalog Number | Dilution | Staining |
|---|---|---|---|---|---|---|
| **APC** | 163Dy | anti-APC | Standard Biotools | 3163001B | 1:50 | Surface |
| **CCL4** | 142Nd | 24006 | R&D | MAB271-100 | 1:100 | Intracellular |
| **CCR7** | 159Tb | G043H7 | Standard Biotools | 3159003A | 1:50 | Surface |
| **CD107a** | 143Nd | H4A3 | BioLegend | 328601 | 1:25 | At the beginning of Stimulation |
| **CD11b (Mac-1)** | 144Nd | ICRF44 | Standard Biotools | 3144001B | 1:50 | Surface |
| **CD11c** | 146Nd | 3,9 | Standard Biotools | 3146014B | 1:50 | Surface |
| **CD123** | 161Dy | 6H6 | BioLegend | 306002 | 1:100 | Surface |
| **CD127 (IL-7Ra)** | 149Sm | A019D5 | Standard Biotools | 3149011B | 1:50 | Surface |
| **CD14** | 151Eu | M5E2 | Standard Biotools | 3151009B | 1:50 | Surface |
| **CD154** | 162Dy | 24-31 | BioLegend | 310802 | 1:50 | Intracellular |
| **CD16** | 209Bi | 3G8 | Standard Biotools | 3209002B | 1:100 | Surface |
| **CD161** | 160Gd | HP-3G10 | BioLegend | 339902 | 1:25 | Surface |
| **CD163** | 154Sm | GHI/61 | Standard Biotools | 3154007B | 1:50 | Surface |
| **CD20** | 147Sm | 2H7 | Standard Biotools | 3147001B | 1:50 | Surface |
| **CD206** | 172Yb | 19,2 | BD Biosciences | 555953 | 1:100 | Surface |
| **CD28** | 175Lu | CD28.2 | BioLegend | 302902 | 1:25 | Surface |
| **CD3** | 170Er | SP34-2 | Standard Biotools | 3170007B | 1:50 | Intracellular |
| **CD4** | 114Cd | L200 | BD Biosciences | 550625 | 1:1000 | Surface |
| **CD45** | 116Cd | D058-1283 | BD Biosciences | 552566 | 1:50 | Surface |
| **CD45RA** | 174Yb | 5H9 | BD Biosciences | 556625 | 1:25 | Surface |
| **CD66abce** | 141Pr | TET2 | Fisher Scientific | MA1-17760 | 1:33 | Surface |
| **CD68** | 106Cd | KP1 | BioLegend | 916104 | 1:25 | Surface |
| **CD69** | 145Nd | FN50 | BioLegend | 310902 | 1:25 | Surface |
| **CD8α** | 111Cd | RPA-T8 | BioLegend | 301053 | 1:50 | Surface |
| **CD8β** | 113Cd | 2ST8.5H7 | Novus Biologicals | NB100-65928 | 1:25 | Surface |
| **CXCR3** | 156Gd | G025H7 | Standard Biotools | 3156004B | 1:50 | Surface |
| **Granzyme B** | 173Yb | GB11 | Standard Biotools | 3173006B | 1:100 | Intracellular |
| **Granzyme K** | 165Ho | GM26E7 | BioLegend | 370502 | 1:50 | Intracellular |
| **HLA-DR** | 176Yb | LN3 | BioLegend | 327002 | 1:400 | Surface |
| **IFN-γ** | 168Er | B27 | Standard Biotools | 3168005B | 1:250 | Intracellular |
| **IgD** | 153Eu | Polyclonal | SouthernBiotech | 2030-01 | 1:25 | Surface |
| **IL-17A** | 148Nd | BL168 | Standard Biotools | 3148008B | 1:100 | Intracellular |
| **IL-1β** | 155Gd | JK1B-1 | BioLegend | 508202 | 1:100 | Intracellular |
| **IL-2** | 158Gd | MQ1-17H12 | Standard Biotools | 3158007B | 1:100 | Intracellular |
| **IL-6** | 167Er | MQ2-6A3 | BD Biosciences | 559068 | 1:100 | Intracellular |
| **MR1 5-OP-RU** | APC | n.a | NIH Tetramer Core | n.a | 1:50 | Surface |
| **NKG2A** | 169Tm | Z199 | Standard Biotools | 3169013B | 1:50 | Surface |
| **Perforin** | 164Dy | Pf-80/164 | Mabtech | 3465-3-250 | 1:100 | Intracellular |
| **TCR Vγ9** | 112Cd | B3 | BioLegend | 331301 | 1:25 | Surface |
| **TCR γδ** | 171Yb | B1 | BioLegend | 331202 | 1:25 | Surface |
| **TNF** | 152Sm | Mab11 | Standard Biotools | 3152002B | 1:500 | Intracellular |
| **DNA Intercalator (Cell ID)** | 191Ir/193Ir | n.a | Standard Biotools | 201192A | 1:100 | Intracellular |
| **Cisplatin (Viability)** | 196Pt | n.a | Standard Biotools | 201064 | 1:10 | Surface |

**Online Table 3. Antibody staining for metal minus multiple experiment.**

| Label | Target | Clone | Tube 1 | Tube 2 | Tube 3 |
|---|---|---|---|---|---|
| **106Cd** | CD68 | KP1 | ✓ | | |
| **111Cd** | CD8α | RPA-T8 | ✓ | | |
| **112Cd** | Vγ9 | B3 | | | |
| **113Cd** | CD8β | SIDI8BEE | | | ✓ |
| **114Cd** | CD4 | L200 | ✓ | | |
| **116Cd** | CD45 | D058-1283 | ✓ | ✓ | ✓ |
| **141Pr** | CD66abce | TET2 | | | ✓ |
| **142Nd** | CCL4 | 24006 | | | |
| **143Nd** | CD107a | 116614 | | | |
| **144Nd** | CD11b | ICRF44 | ✓ | | |
| **145Nd** | CD69 | FN50 | | ✓ | |
| **146Nd** | CD11c | 3.9 | | | ✓ |
| **147Sm** | CD20 | 2H7 | ✓ | | |
| **148Nd** | IL-17A | BL168 | | | |
| **149Sm** | CD127 | A019D5 | | ✓ | |
| **151Eu** | CD14 | M5E2 | | | ✓ |
| **152Sm** | TNF | Mab11 | | | |
| **153Eu** | IgD | Polyclonal | ✓ | | |
| **154Sm** | CD163 | GHI/61 | | ✓ | |
| **155Gd** | IL-1β | JK1B-1 | | | |
| **156Gd** | CXCR3 | G025H7 | | | ✓ |
| **158Gd** | IL-2 | MQ1-17H12 | | | |
| **159Tb** | CCR7 | G043H7 | ✓ | | |
| **160Gd** | CD161 | HP-3G10 | | ✓ | |
| **161Dy** | CD123 | 6H6 | | | ✓ |
| **162Dy** | CD154 | 24-31 | | | |
| **163Dy** | MR1 | anti-APC | | ✓ | |
| **164Dy** | Perforin | Pf-80/164 | ✓ | | |
| **165Ho** | Granzyme K | GM26E7 | | | ✓ |
| **167Er** | IL-6 | MQ2-6A3 | | | |
| **168Er** | IFN-γ | B27 | | | |
| **169Tm** | NKG2A | Z199 | | ✓ | |
| **170Er** | CD3 | SP34-2 | ✓ | | |
| **171Yb** | Pan TCR γδ | B1.1 | | | ✓ |
| **172Yb** | CD206 | 19.2 | | ✓ | |
| **173Yb** | Granzyme B | GB11 | ✓ | | |
| **174Yb** | CD45RA | 5H9 | | | ✓ |
| **175Lu** | CD28 | CD28.2 | | ✓ | |
| **176Yb** | HLA-DR | LN3 | ✓ | | |
| **209Bi** | CD16 | 3G8 | | | ✓ |

### ONLINE FIGURES

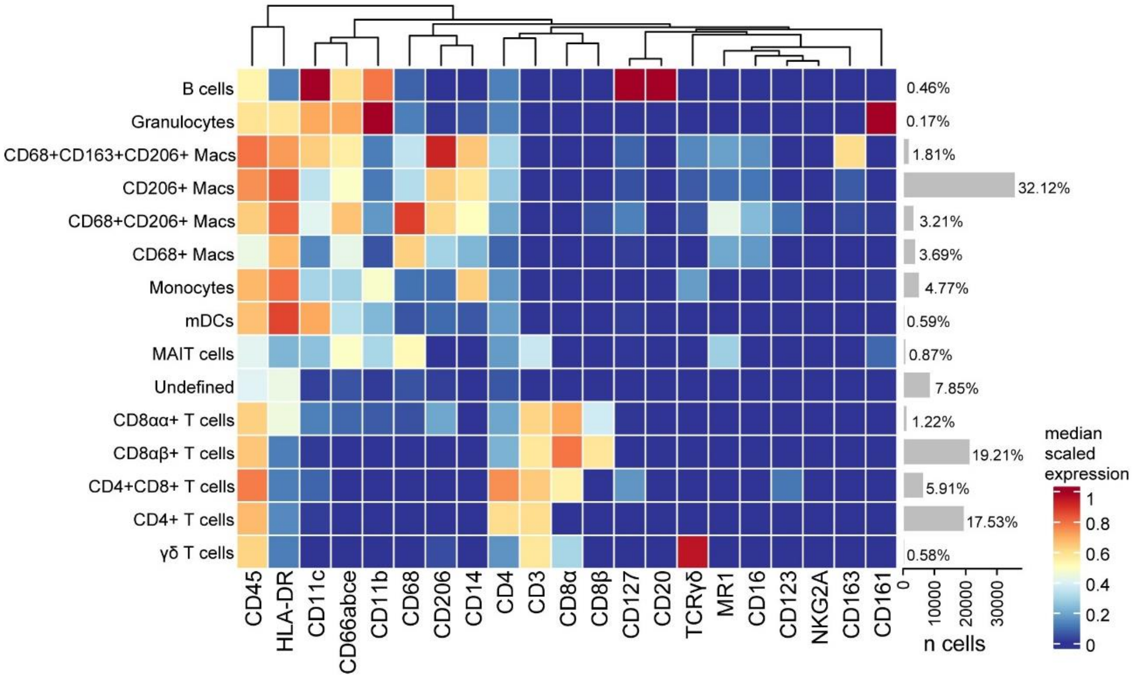


**Online Figure 1:** Identification of leukocytes subsets in BAL using FlowSOM clustering. Live CD45+ leukocytes were exported from FlowJo and FlowSOM clustering algorithm was used to perform unsupervised clustering based using canonical makers. FlowSOM clusters were then manually annotated based on the expression of canonical markers as identified using FlowJo. However, FlowSOM could not identify the NK subset without overclustering.

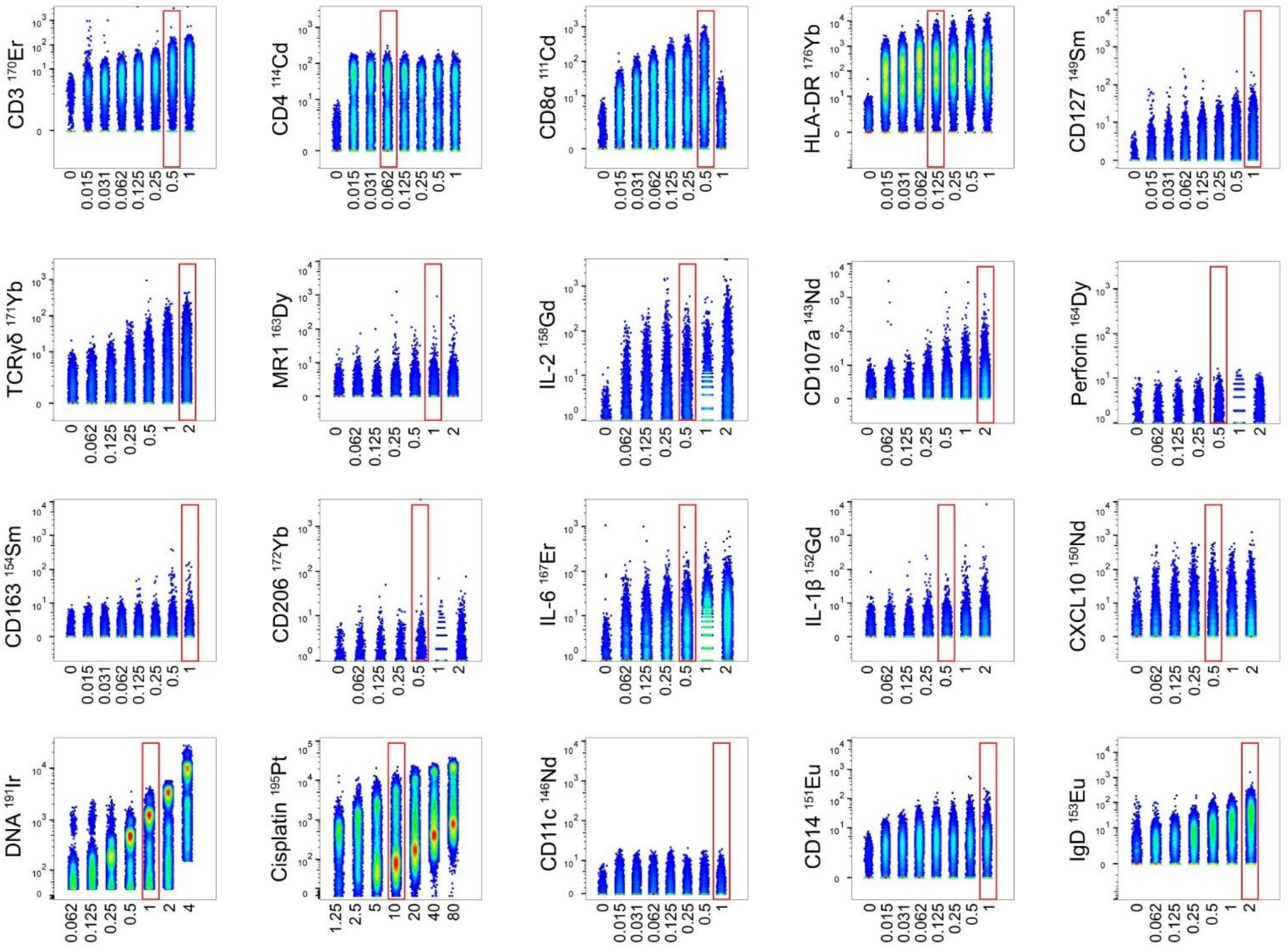


**Online Figure 2:** Serial titrations. Two-fold serial dilutions of reagents were tested on cryopreserved splenocytes from rhesus macaques. The total staining volume for each titration was 50 µL, except for DNA intercalator staining which was performed in a total of 100 µL. Files of each titration experiment were concatenated. Red rectangles indicate the chosen titer based on the frequencies of positive cells and spillover into mass+/-1 and oxidation (+16) channels. Final titration results are presented in Online Table 2.

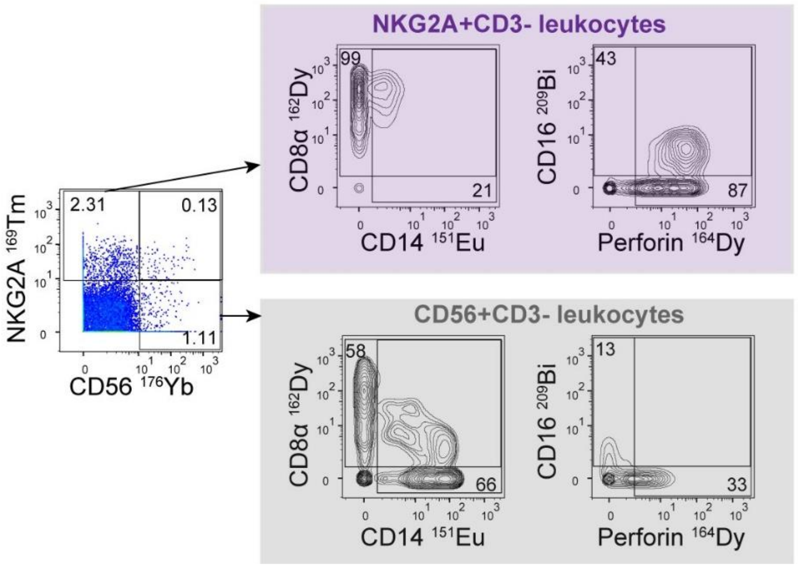


**Online Figure 3:** Comparison of NKG2A and CD56 markers for identification of NK cells on splenocytes from rhesus macaques. Co-staining of NKG2A and CD56 indicated that the majority of cells stained single positive for NKG2A or CD56 with twice the frequency of NKG2A+ cells compared to CD56. NKG2A positive cells almost exclusively CD8α+ and expressed a higher percentage of CD16 and perforin compared to CD56. In contrast, CD56 positive cells expressed a higher percentage of CD14 compared to NKG2A positive cells.

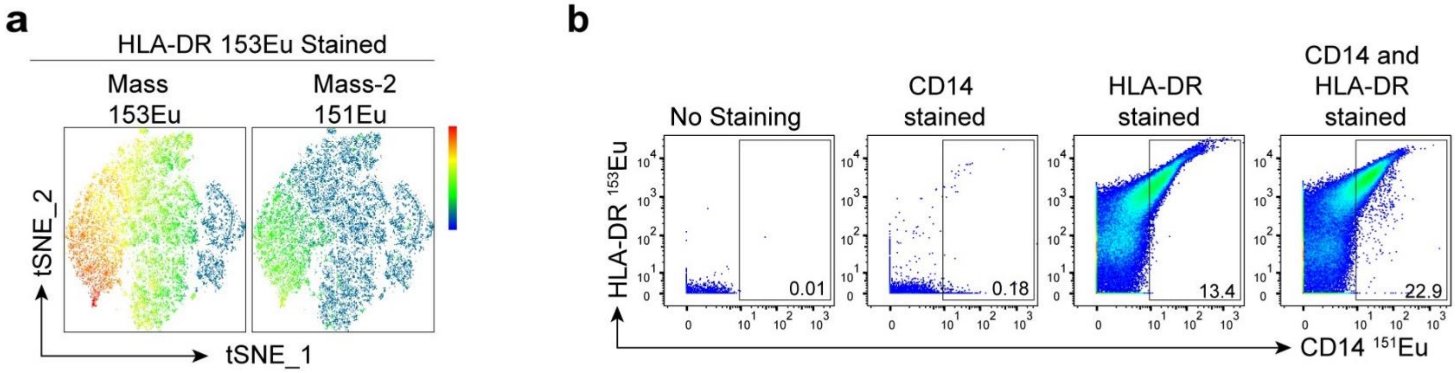


**Online Figure 4:** Examination of HLA-DR 153Eu spillover into CD14 151Eu. **(A)** tSNE dimensionality reductions plots showing signal in the mass-2 (CD14 151Eu channel) when cells were stained with HLA-DR 153Eu and not CD14 151Eu. The observed spillover signal corresponds to the highest HLA-DR scaled expression signal intensity. **(B)** Bivariate plots showing the detection of CD14 151Eu positive events when CD14 was stained in the absence vs the presence of HLA-DR staining and when HLA-DR 153Eu was stained in the absence vs the presence of CD14 staining. Unstained condition was included as a negative control.

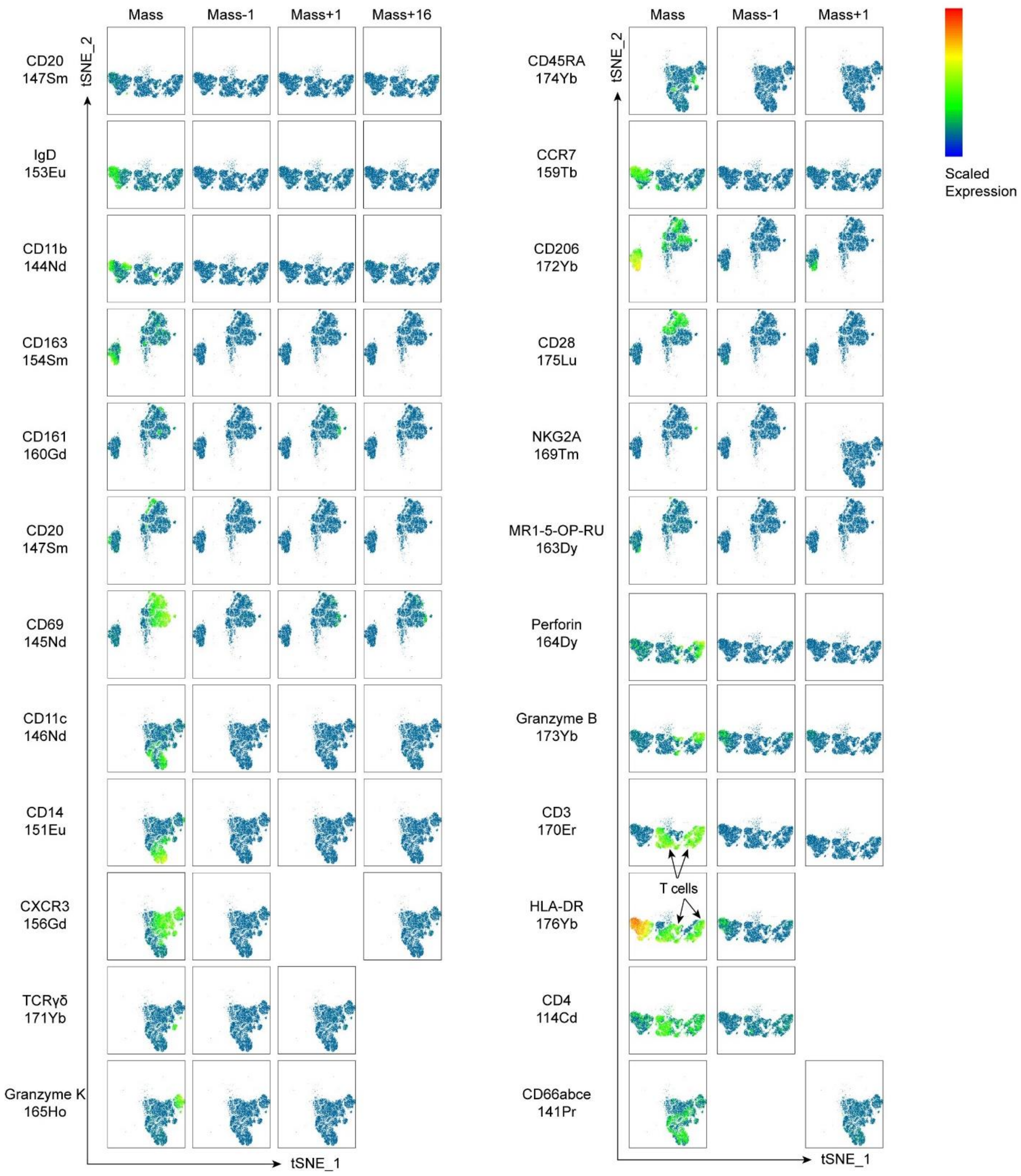


**Online Figure 5:** Examination of spillover using tSNE dimensionality reductions plots showing scaled median marker expression. A metal minus multiple experiment was performed for antibodies that did not require stimulation. Staining was divided into three tubes making sure to leave open channels in mass-1, mass+1 and mass+16 for spillover assessment. Metal impurity/abundance sensitivity spillover of CD206 Yb172 signal into granzyme B Yb173 was detected, but this does not affect data analysis because granzyme B is characterized on CD206 negative cells. We also observed signal spillover from HLA-DR 176Yb into CD28 175Lu (mass-1) due to abundance sensitivity. The location of the HLA-DR 176Yb spillover signal on the tSNE plot associated with the channels for CD28 175Lu suggests the spillover signal was limited to non-T cells (CD3-), and therefore does not affect CD28 expression analysis profiled on T cells. Arrows indicate the location of T cells on the tSNE plots.
